# Toggling of NKG2A expression drives functional specialization of iPSC-derived CAR NK cells

**DOI:** 10.1101/2025.08.20.671199

**Authors:** Minoru Kanaya, Camille Philippon, Herman Netskar, Michelle Lu Saetersmoen, Artur Cieslar-Pobuda, Lamberto Torralba-Raga, Giovanna Perinetti Casoni, Quirin Hammer, Marianna Vincenti, Merete Thune Wiiger, Silje Zandstra Krokeide, Hanna Julie Hoel, Eivind Heggernes Ask, Mizuha Kosugi-Kanaya, Lise Kveberg, Hui-yi Chu, Brian Groff, Jeffrey S. Miller, Tom Lee, Dan S Kaufman, Jode P Goodridge, Bahram Valamehr, Aline Pfefferle, Karl-Johan Malmberg

## Abstract

Induced pluripotent stem cell (iPSC)-derived natural killer (iNK) cells offer a promising platform for off-the-shelf immunotherapy against hematological malignancies. NK cell function is dynamically regulated through education driven by inhibitory receptors, including CD94/NKG2A and killer cell immunoglobulin-like receptors (KIR). However, the acquisition of inhibitory receptors in iNK cells and their role during differentiation and education remains poorly defined. In this study, we monitored receptor repertoires, transcriptional states, and functional responses in a range of genetically engineered iNK cell lines. Transcriptional reference mapping placed iNK cells close to cytokine-activated NKG2A^+^ CD56^dim^ peripheral blood (PB) NK cells. Despite their early differentiation stage, iNK cells displayed a well-developed cytotoxic effector program, which was also reflected in high protein expression of Eomes, granzyme B, and activating receptors DNAM-1 and NKG2D. Acquisition of NKG2A by iNK cells was associated with a more differentiated transcriptional state and superior functional responses against a broad range of targets, including those expressing low to moderate levels of HLA-E, suggesting attenuated inhibitory signaling through NKG2A in iNKs. CRISPR knockout of β2-microglobulin (*Β2μ*) in iNK cells revealed that the functional potency of NKG2A^+^ iNK cells was independent of educating interactions with HLA-E in cis or trans. Finally, CRISPR-mediated ablation of NKG2A led to a spontaneous compensatory surface expression of CD94/NKG2C heterodimers, associated with enhanced IFN-*γ* production and cytotoxic activity against target cells with forced high expression of single-chain β2m-HLA-E-peptide trimers. Our results indicate an education-independent functional maturation of iNK cells, characterized by potent effector programs coupled with a favorable early-stage transcriptional profile.

## Introduction

Differentiation of human induced pluripotent stem cells (iPSCs) into functional immune effector cells is a promising approach to generate a broad array of off-the-shelf cell therapies against cancer and other disorders.^1, 2^ The main benefits of the iPSC platform are the possibility to perform multiplexed gene editing at the clonal level and to scale manufacturing for high accessibility on demand.^1^ Genetically engineered iPSC-derived natural killer (iNK) cells show great efficacy in several experimental tumor models, including acute myeloid leukemia (AML), multiple myeloma (MM) and B cell lymphoma but also against solid tumors in combination with NK cell specific engagers.^3, 4, 5, 6, 7, 8, 9, 10, 11, 12^ Off-the-shelf iNK cells have also reached early Phase I clinical trials and demonstrated a favorable safety profile with low risk of graft-versus-host disease or cytokine release syndrome (CRS), and some anti-tumor activity in patients with advance stage disease.^13^

The promising pre-clinical and clinical data of the first-generation iNK cell products has sparked an interest in designing synthetic cells with enhanced intrinsic functional properties combined with smart receptor logic for controlled activation in complex heterogenous tumor environments.^1, 14^ A necessary requirement to achieve this goal, is to understand how the functional template evolves in iNK cells to identify steps during which engineering can further boost the functional template. Under homeostatic conditions, NK cell function develops during a transcriptionally regulated differentiation process from CD56^bright^ NK cells (also termed NK2) via early NKG2A^+^ CD56^dim^ NK cells (NK1a) to intermediate and late CD56^dim^ stages (NK1b-c) with acquisition of killer cell immunoglobulin-like receptors (KIR), CD57 and a gradual loss of NKG2A.^15, 16, 17^ In some individuals, infection with cytomegalovirus (CMV) drives the differentiation of adaptive NKG2C^+^ NK cells (NK3), representing a more terminal transcriptional state, associated with profound epigenetic remodeling.^18, 19, 20^ During terminal differentiation, NK cells become gradually less responsive to cytokines but acquire a strong cytotoxic function upon the engagement of activating receptors including the FcγRIII (CD16) and NKG2C in adaptive NK cells.^21, 22^

In parallel with differentiation, NK cells are functionally fine-tuned during a process termed education, driven by inhibitory receptors binding to self HLA class I.^17, 21, 23, 24^ These interactions shape the missing-self response of NK cells like a rheostat, e.g., the stronger the inhibitory input is during education, the stronger the functional response to target cells lacking the cognate ligand.^25^ There are two main “schools” of education, one determined by interactions of CD94/NKG2A with HLA-E and one by inhibitory killer cell immunoglobulin-like receptors (KIR) binding to different HLA-A, B and -C motifs.^26^ Surprisingly, there is no distinct gene signature associated with NK cell education.^23, 24^ Nevertheless, there are several phenotypic traits associated with the educated state, which provide some clues regarding the underlying mechanism. Educated NK cells express high levels of DNAM-1^27, 28^ and display increased intracellular stores of effector molecules granzyme B and perforin.^23^ They also have an altered metabolism with increased mTOR activity and glycolysis,^29, 30^ unique receptor compartmentalization in the cell membrane with decreased densities of self-HLA specific KIRs and the associated SHP1 molecules at the immune synapse.^31, 32, 33^ How these phenotypes are connected remains unclear, but it seems plausible that the cell-cell interactions during which inhibitory receptors interact with their cognate ligand, shape the interior of the cell and give rise to many, if not all, of these phenotypes that determine the functionality.

In this study we examined the transcriptional and functional state of engineered iNK cells by transfer learning and mapping onto a reference of resting and IL-15 stimulated peripheral blood (PB) NK cell subsets. We found that feeder-expanded iNK cells have a differentiation stage close to cytokine stimulated early NKG2A^+^ CD56^dim^ NK cells. Introduction of tonic IL-15/IL-15R signaling combined with chimeric antigen receptor (CAR) expression was associated with increased differentiation, which was also reflected in acquisition of NKG2A and KIR receptors. Subset stratification and CRISPR/Cas9 deletion of *B2M* showed that the highly functional profile of NKG2A^+^ iNK cells was independent of education and relatively insensitive to physiological levels of the HLA-E checkpoint. CRISPR-based knockout of *KLRC1* led to a spontaneous surface expression of NKG2C and cooperative signaling between the CAR and NKG2C against CD19^+^ Nalm-6 with forced overexpression of HLA-E. These results shed light on the regulatory programs involved in the functional development and diversification of the iNK cell template and may inform future designs of off-the-shelf cell therapy.

## Results

### Transcriptional reference mapping of iPSC-derived NK cells

We generated a series of iNK cell lines and performed genetic engineering as schematically described in **Figure 1A** and **Supplementary Figure 1A**. Cellular indexing of transcriptomes and epitopes by sequencing (CITE-Seq) analysis of unedited and CAR19/IL-15/IL-15R engineered iNK cells was performed and the transcriptomes were mapped onto a reference of major peripheral blood (PB)-derived human leukocytes, including CD4 and CD8 T cells, CD56^bright^ and CD56^dim^ NK cells, NKT cells, ψ8T cells, B cells and myeloid cells. iNK cells were annotated as NK cells by CellTypist (**Supplemental Figure 1B**)^34^. We applied Partition-based graph abstraction (PAGA)^35^ to quantify the connectivity of iNK cell subsets with the PB-NK cell subsets in the reference (**Figure 1B-C**). Reference mapping of iNK cells and IL-15 stimulated NK cell subsets placed both unedited and engineered iNK cells closest to rapidly cycling intermediate-stage CD56^dim^ NK cells (**Figure 1D-E and Supplementary Figure 1B**).

**Figure 1.**
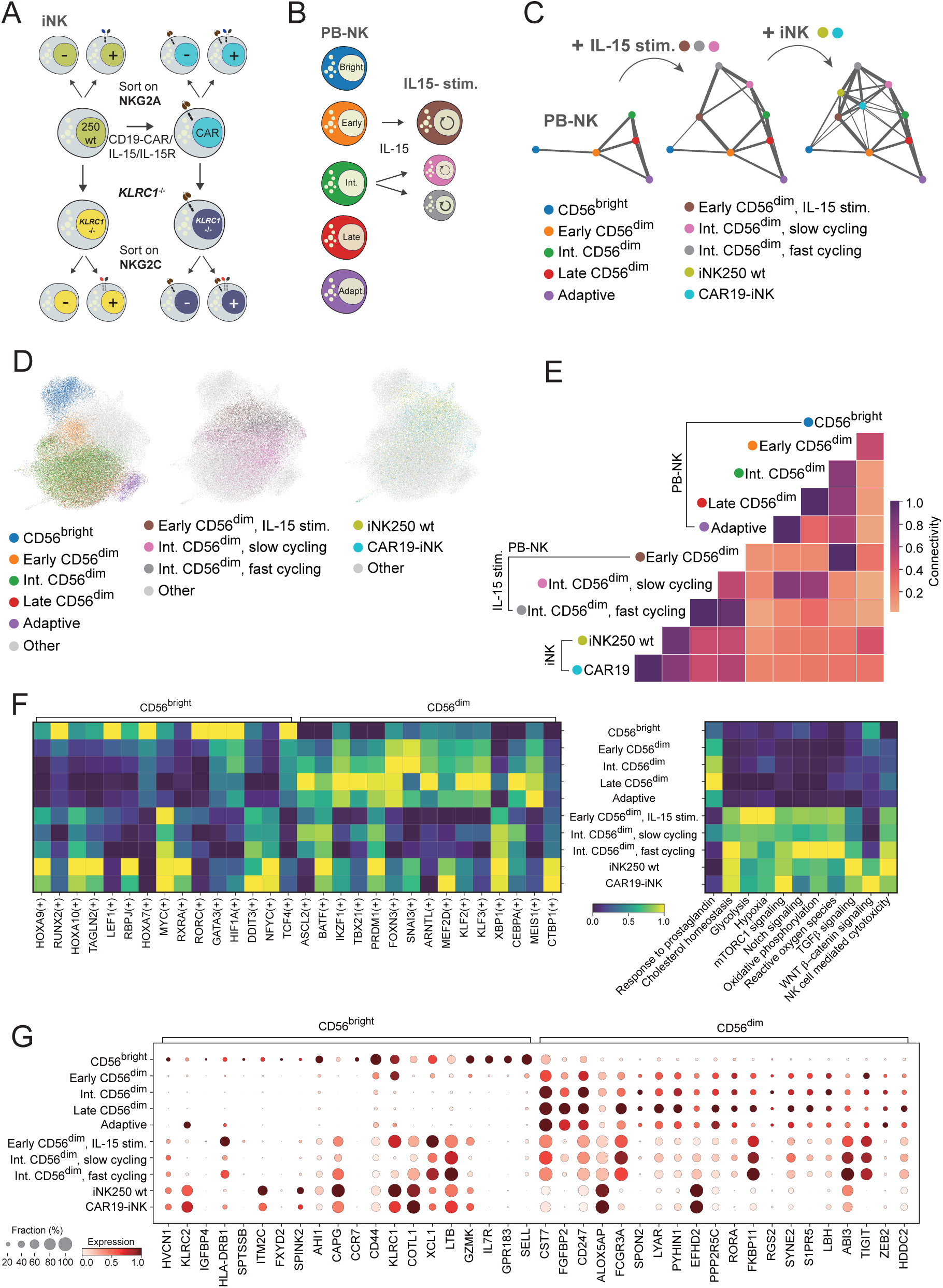
Transcriptional reference mapping of iPSC-derived NK cells. (**A**) Schematic of iNK cell generation and genetic engineering strategies, including CD19 CAR/IL-15/IL-15R transduction and *KLRC1* knockout. (**B**) Schematic overview of peripheral blood NK (PB-NK) and IL-15 stimulated NK cell single-cell RNA sequencing datasets. (**C-D**) Mapping of iNK cells onto a PB-NK reference. (**E**) Partition-based graph abstraction (PAGA) depicting connectivity between engineered iNK subsets and PB-NK cells. (**F-G**) Transcriptional states, regulatory programs and comparison of differentiation sages of iNK subsets compared to PB-NK subsets. n = 2-12.

Next, we constructed gene regulatory networks (GRN)^36^ and compared the states of iNK subsets relative to the dominant regulons defining stages of normal NK cell differentiation.^16^ This analysis corroborated the intermediate maturation stage of iNK cells with high expression of several programs associated with both resting CD56^bright^ and CD56^dim^ regulons but with several distinct programs being either turned on or off. While most of these, including MYC and LEF were shared with IL-15 stimulated PB-NK cells, iNK cells also displayed some unique features, including high WNT signaling (**Figure 1F**). Both IL-15 stimulated PB-NK cells and wt/CAR iNK cells displayed upregulation of several functional programs, including cholesterol homeostasis, Notch signaling, glycolysis, oxidative phosphorylation, mTOR signaling and transcriptional programs associated with cytotoxicity. The only downregulated program identified was response to prostaglandins. Among hallmark genes of CD56^bright/dim^ NK cells, iNK cells expressed high levels of both *KLRC1* and *KLRC2*, encoding for NKG2A and NKG2C, respectively (**Figure 1G**). iNK cells expressed very high levels of *EFHD2*, a positive regulator of T cell receptor signaling,^37, 38^ and *ALOX5AP*, involved in the immunological circuits of the inflammatory response.^39^ iNK cells were uniquely low in *CST7, FGFBP2, SPON2, LYAR*, *SYNE2, TIGIT* and *FCGR3A* (**Figure 1G**). *CST7* is a negative regulator of Cathepsin C which cleaves pro-granzyme B to the active form of granzyme B.^13^ Together these data demonstrate efficient iPSC differentiation into bona fide NK cells with a transcriptional profile that bears striking similarities with IL-15 stimulated primary PB-derived NK cells.

### NKG2A expression defines functionally mature iNK cells

Given the high level of transcripts of both *KLRC1* and *KLRC2* in iNK cells we monitored surface expression of NKG2A and NKG2C in two distinct wildtype (wt) iNK cell lines (121 and 250) and iNK250 cells engineered to express a previously described CAR19 construct containing the transmembrane domain of NKG2D, the 2B4 co-stimulatory domain, and the CD3zeta signaling domain as well as an IL-15/IL-15R fusion protein (CAR19-iNK).^7^ Expansion of PB-NK with K562-41BB-mIL21 feeder cells led to higher frequencies of NKG2A^+^ NK cells. NKG2A expression frequencies increased further in iNK cells transduced with CAR and IL-15/IL-15R, providing tonic cytokine stimulation (**Figure 2A**). In addition to NKG2A, a small subset of NKG2C^+^NKG2A^+^ double positive iNK cells was present in CAR19-iNK cells. In line with previous studies, both resting NKG2A^+^ PB-NK cells and expanded NKG2A^+^ PB-NK cells demonstrated significantly higher degranulation and IFN-*γ* production against K562 cells compared to NKG2A^-^ NK cells^40, 41, 42^ (**Figure 2B**). We found that NKG2A^+^ iNK121 wt cells, iNK250 wt cells and CAR19-iNK cells showed higher responsiveness compared to NKG2A^-^ iNK cells, both in natural cytotoxicity assays against K562 (**Figure 2C-D**) and for CAR19-iNK cells also against CD19^+^ Nalm-6 cells (**Figure 2E**). The increased degranulation responses of NKG2A^+^ iNK cells were corroborated in Incucyte-based killing assays against K562 using FACS sorted NKG2A^+^ and NKG2A^-^ iNK250 cells (**Figure 2F**).

**Figure 2.**
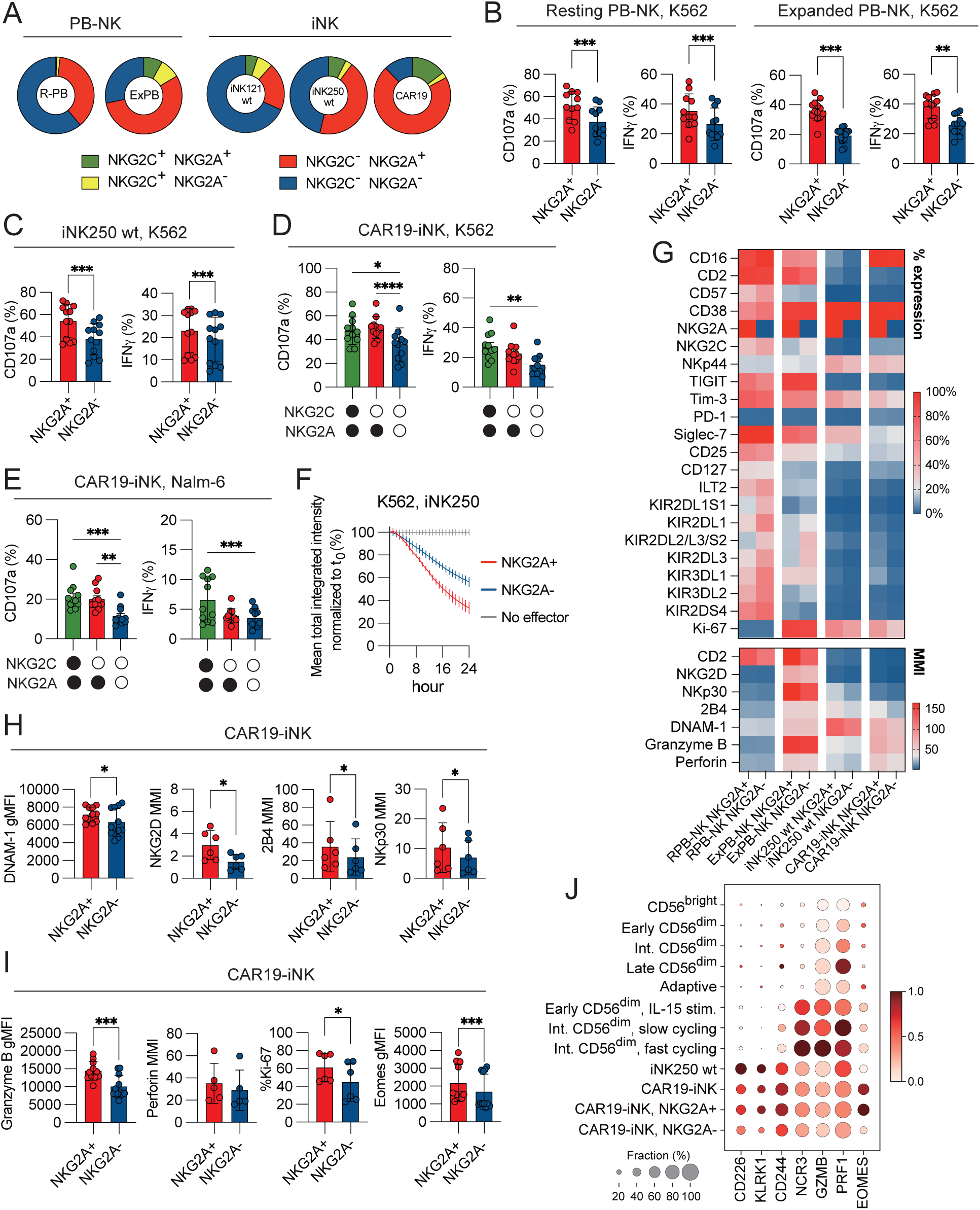
Functional superiority of NKG2A+ iNK cells. (**A**) Flow cytometric analysis of NKG2A and NKG2C expression in resting and expanded PB-NK cells and iNK lines (iNK121 wt, iNK250 wt, and CAR19-iNK). (**B-E**) Degranulation (CD107a) and IFN-γ production assays demonstrate enhanced cytotoxicity of NKG2A+ subsets compared to NKG2A-subsets in PB-NK and iNK lines against K562 (**B-D**) and Nalm-6 targets (**E**). (**F**) Cytotoxicity of NKG2A+ compared to NKG2A-iNK cells against K562 target cells. (**G**) Phenotypic characterization of resting PB-NK, expanded PB-NK and iNK lines (iNK205, CAR19-iNK) stratified by NKG2A expression. (**H-J**) Granzyme B, perforin, DNAM-1, and Eomes expression levels in NKG2A+ CAR19-iNK cells.. Data are represented as mean (SD). Significance was calculated using a Wilcoxon matched-pairs signed rank test (**B-C, H-I**) or a Friedman test followed by Dunn’s multiple comparison test (**D-E**). p values: * < 0.05, ** < 0.01, *** < 0.001, **** < 0.0001. n = 2-13.

Next, we used mass cytometry and flow cytometry to perform an extensive phenotypic analysis of NK cell receptors, transcription factors and effector molecules in NKG2A^+^ or NKG2A^-^ resting and expanded PB-NK, wt and CAR engineered iNK cell lines (**Figure 2G**). We confirmed an intermediate maturity of the iNK cells with low expression of typical markers of maturity, including CD16, KIR, CD57 and NKG2C. Notably, CD16 expression was restored in the CAR19-iNK line carrying the non-cleavable high-affinity CD16 transgene.^3^ Although KIR expression was generally low, we found that the CAR19-iNK cell variant carrying the IL-15/IL-15R construct displayed higher levels of KIR (**Supplementary Figure 2A**). Stratification of CAR19-iNK cells based on NKG2A expression revealed an inverted pattern of KIR and NKG2A co-expression compared to PB-NK cells which exhibited higher KIR expression on NKG2A^+^ iNK cells (**Supplementary Figure 2B**), suggesting NKG2A acquisition reflects a late stage of differentiation in iNK cells. We also found elevated levels of several activating receptors, including DNAM-1, NKG2D, and NKp30, as well as cytotoxic granules containing granzyme B and perforin (**Figure 2G-I**). Transcriptionally, NKG2A^+^ CAR19-iNK cells exhibited higher levels of Eomes, a master regulator of NK cell function, compared to NKG2A^-^ iNK cells (**Figure 2J)**. These data establish that NKG2A expression correlates with a mature effector phenotype in iNK cells, characterized by enhanced cytotoxicity, cytokine production, and expression of activating receptors.

### The functional maturation of NKG2A^+^ iNK cells is independent of HLA-E driven education

Thus far, we established that NKG2A expression reflects transcriptional, phenotypic and functional maturity in iNK cells. To determine whether the functional superiority of NKG2A^+^ iNK cells depends on HLA-E-mediated education,^26^ we generated two independent β2-microglobulin knockout (iNK *B2M*^-/-^) lines (**Figure 3A-C**). Flow cytometric analysis confirmed significantly reduced levels of HLA-ABC and HLA-E on *B2M*^-/-^ iNK cells compared to wild-type iNK cells (**Figure 3B**). Despite the absence of HLA-E, NKG2A^+^ *B2M*^-/-^ iNK cells retained their functional advantages over NKG2A^-^ iNK cells (**Figure 3D-E**). These findings were consistent across multiple iNK lines, confirming that NKG2A expression in iNK cells is associated with functional maturation independent of classical education via HLA-E interactions. Therefore, we propose that this mechanism represents a distinct regulatory pathway for iNK cell functionality, divergent from PB-NK cells.

**Figure 3.**
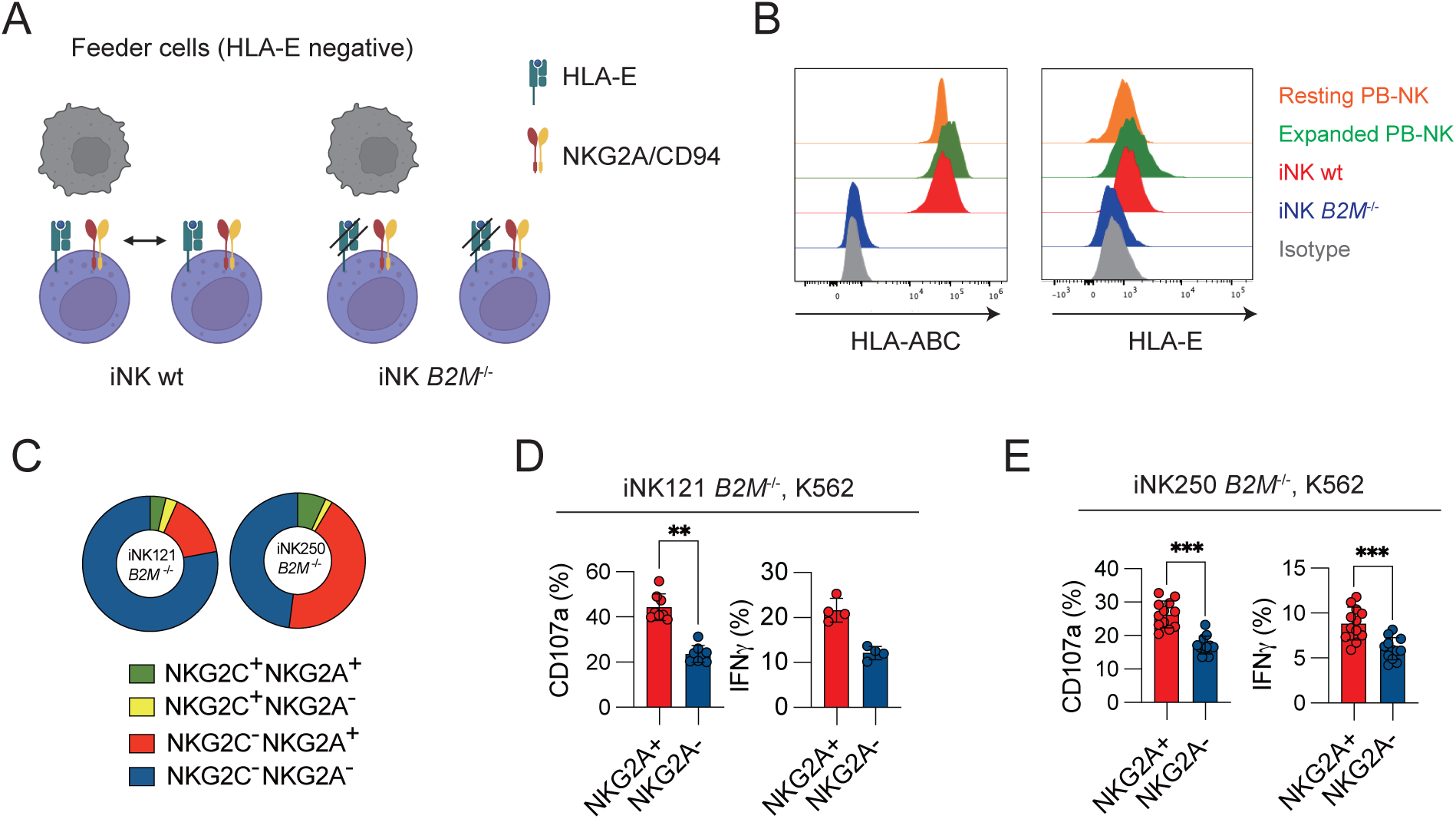
Education-independent functional maturation of NKG2A+ iNK cells. (**A-B**) Schematic overview and surface expression of HLA-ABC and HLA-E in wild-type (iNK wt) and β2-microglobulin-deficient (iNK *B2M*^-/-^) iNK cells. (**C**) Distribution of NKG2A and NKG2C subsets in the iNK121 *B2M*^-/-^ and iNK250 *B2M*^-/-^ cell lines. (**D-E**) Functional assays (CD107a, IFNγ) of iNK121 *B2M*^-/-^ and iNK250 *B2M*^-/-^ cells against K562 target cells, stratified by NKG2A expression. Data are represented as mean (SD). Significance was calculated using a Wilcoxon matched-pairs signed rank test. p values: * < 0.05, ** < 0.01, *** < 0.001, **** < 0.0001. n = 4-12.

### Attenuated inhibition through the HLA-E-NKG2A inhibitory axis in iNK cells

We investigated the impact of varying HLA-E expression levels on iNK cell functionality by utilizing target cell lines with differential HLA-E expression, including K562 and Nalm-6 variants. Flow cytometric analysis confirmed graded HLA-E levels on the engineered target cells, ranging from negative (Nalm-6 HLA-E^-/-^), low (K562), to physiological (K562 HLA-E, Nalm-6) and supraphysiological expression (K562/Nalm-6 HLA-E^high^) (**Figure 4A-B**). Degranulation and IFN-*γ* responses of PB-NK cells, wt iNK cells and CAR19-iNK cells were largely unaffected by low to intermediated levels of HLA-E on target cells but significantly dampened by high levels of HLA-E (**Figure 4C-D** and **Supplementary Figure 3A**-E). This effect on natural cytotoxicity against K562 variants was subset specific and only seen in NKG2A^+^ iNK cells. In NKG2A^+^NKG2C^+^ double positive iNK cells, the inhibitory signaling dominated, in line with the higher affinity of the NKG2A receptor (**Figure 4D**)^43, 44, 45^. Similar functional response profiles were observed in resting and expanded PB-NK cells (**Supplementary Figure 3A**-E). Wildtype levels of HLA-E in Nalm-6 cells was only marginally inhibitory as revealed by slightly increased functional responses of CAR19-iNK cells against HLA-E^-/-^ Nalm-6 cells (**Figure 4E**). However, overexpression of HLA-E led to significantly reduced responses also by CAR engineered iPSCs (**Figure 4E**), an effect that was more pronounced in NKG2A^+^ CAR19-iNK cells (**Figure 4F-G**). These results underscore the importance of addressing HLA-E-mediated inhibition in CAR19-iNK cell therapies, particularly in the context of malignancies with elevated HLA-E expression.

**Figure 4.**
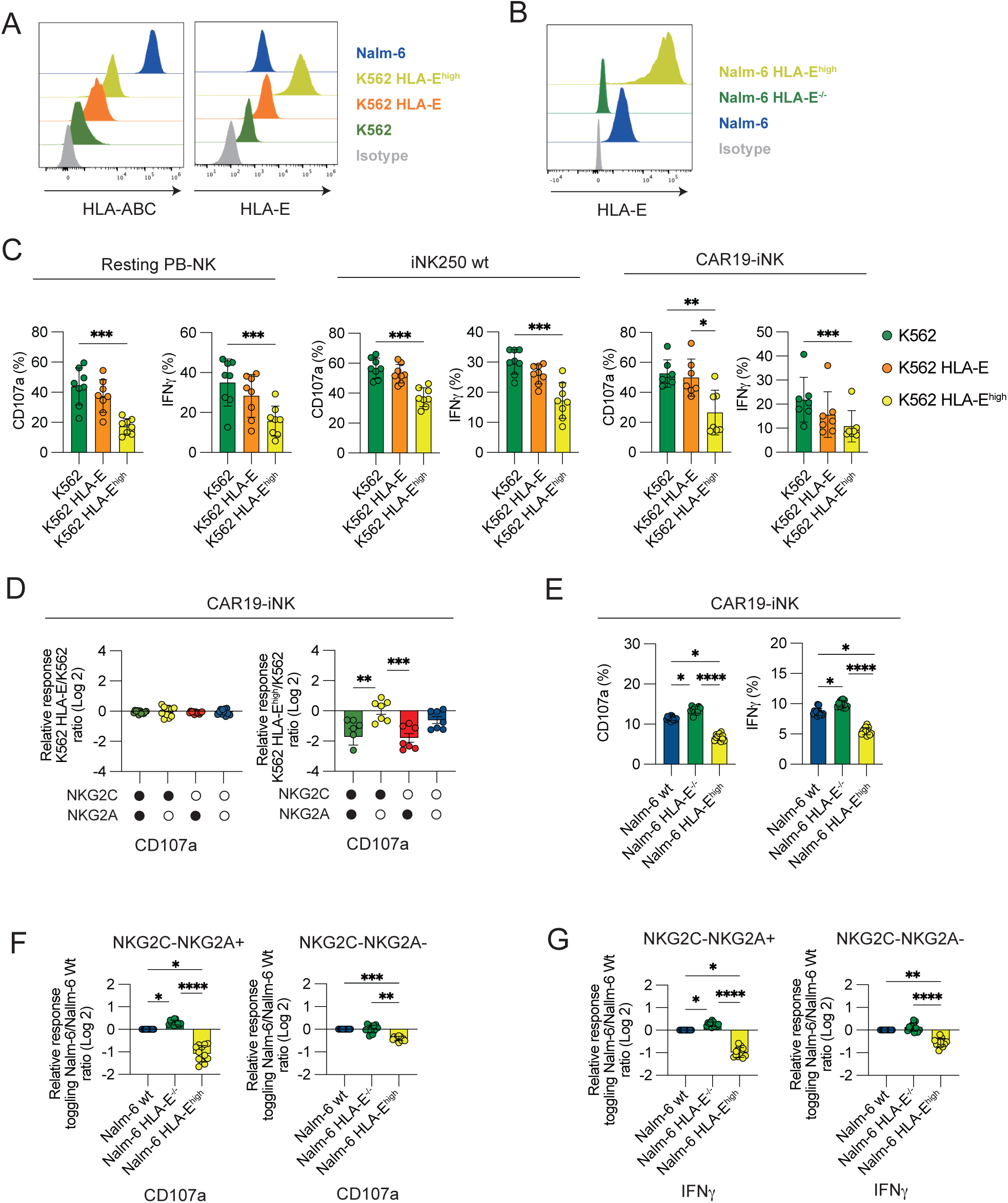
HLA-E levels modulate iNK cell functionality. (**A-B**) Flow cytometric analysis of HLA-ABC and HLA-E expression in target cells, including K562 and Nalm-6 cells with varying HLA-E expression levels. (**C**) Degranulation (CD107a) and IFNγ assays demonstrate of resting PB-NK, iNK250 and iNK-CAR19 cells to targets with supraphysiological HLA-E expression. (**D**) Relative response rate calculations of NKG2A+ and NKG2A-subsets against targets with varying levels of HLA-E expression (**E**) Degranulation (CD107a) and IFNγ production of CAR19-iNK against Nalm-6 cell lines expressing varying HLA-E levels. (**F-G**) Functional responses of CAR19-iNK cells expressing or lacking NKG2A against Nalm-6 HLA-E variants. Data are represented as mean (SD). Significance was calculated using a Friedman test followed by Dunn’s multiple comparison test. p values: * < 0.05, ** < 0.01, *** < 0.001, **** < 0.0001. n = 7-12.

### Genetic ablation of *KLRC1* (NKG2A) leads to compensatory surfacing of NKG2C

Given our observation that the functional responses of the NKG2A^+^ iNK cell subset were independent of educating interactions with HLA-E in cis or trans, we speculated that genetic deletion of *KLRC1* would have no negative impact on the function and instead could unleash more potent responses against target cells expressing high levels of HLA-E. CRISPR-based deletion of *KLRC1* in CAR19-iNK cells (CAR19-iNK *KLRC1*^-/-^) was carried out at the iPSC stage and was confirmed in iNK cells derived from these clones (**Figure 5A**). Surprisingly, genetic ablation of *KLRC1* in CAR19-iNK led to distinct surface expression of NKG2C (**Figure 5A-B**). This was observed in two independent lines (**Supplementary Figure 4A)** and was not linked to increased *KLRD1* (CD94) or *KLRC2* (NKG2C) transcripts as determined by CITE-seq (**Figure 5B-C**). Notably, knockout of *KLRC1* in PB-NK cells did not lead to a similar surface expression of NKG2C which remained restricted to the adaptive NK cell subset (**Supplementary Figure 4B**-C). We found that baseline levels of *KLRC2* were higher in both wt and *KLRC1^-/-^* iNK cells compared to PB-NK cells, reaching similar levels to those observed in adaptive NK cells (**Figure 5C**). Furthermore, despite slightly reduced expression levels of CD94 in CAR19-iNK *KLRC1*^-/-^, heterodimerization of NKG2C and CD94 was confirmed in FACS analysis (**Supplemental Figure 4D**). Thus, we observed a unique compensatory surface expression of NKG2C in *KLRC1*^/-^ iNK cells with high baseline levels of *KLRC2* transcripts.

**Figure 5.**
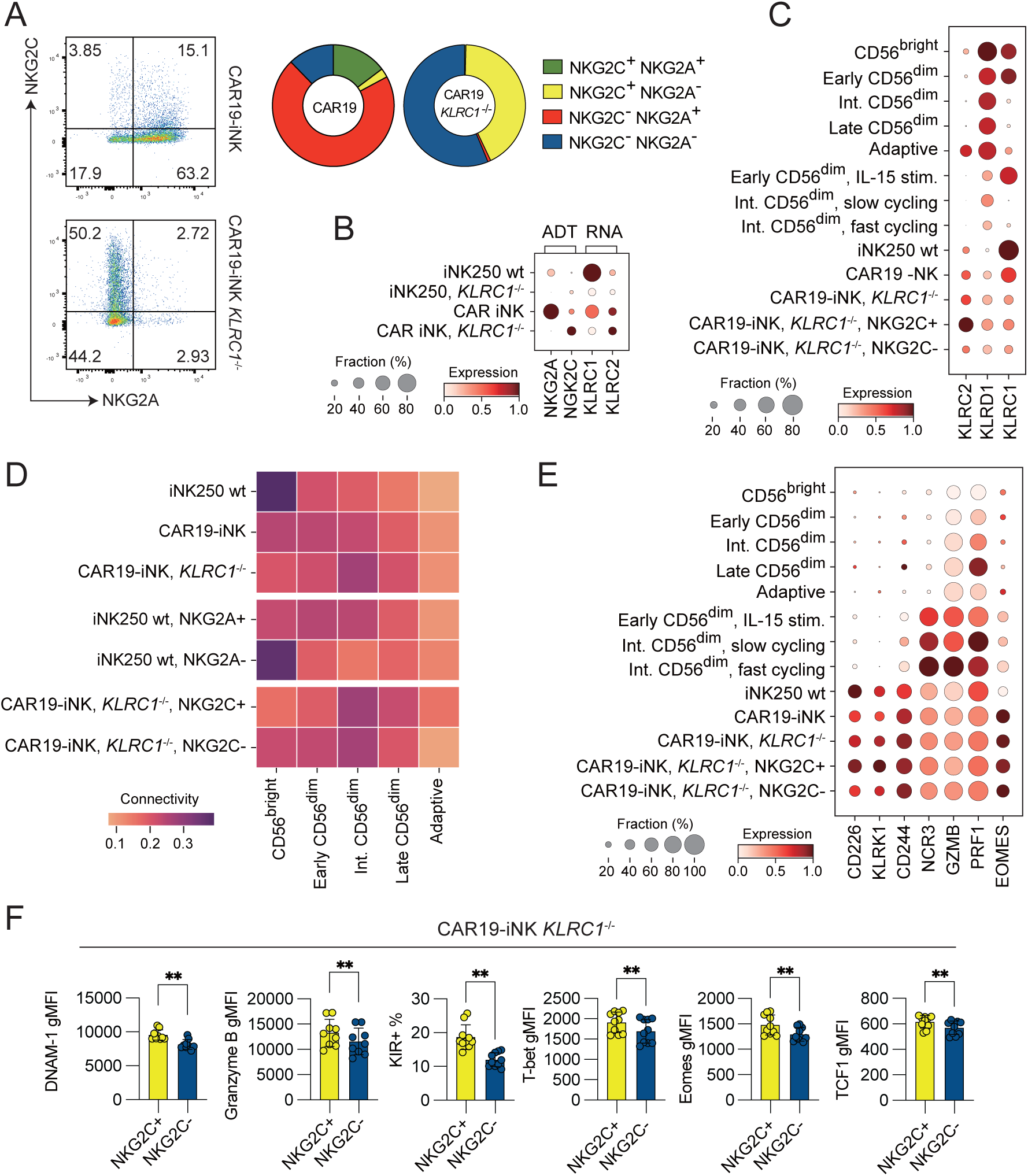
Adaptive reprogramming of CAR19-iNK cells following *KLRC1* knockout. (**A-C**) NKG2C expression at the protein (**A-B**) and transcript level (**B-C**) after CRISPR knockout of *KLRC1* in iNK-CAR19 cells. (**C-D**) Mapping of iNK cell lines onto a PB-NK reference map. (**E-F**) Expression of activating receptors, effector molecules, and transcription factors in reference populations and CAR19-iNK KLRC1^-/-^ cells stratified by NKG2C expression. Data are represented as mean (SD). Significance was calculated using a Wilcoxon matched-pairs signed rank test. p values: * < 0.05, ** < 0.01, *** < 0.001, **** < 0.0001. n = 2-12.

We next performed reference mapping of iNK cell variants, including wt-iNK, CAR19-iNK cells, NKG2A+ or NKG2A-CAR-iNK cells, bulk *KLRC1*^-/-^ CAR-iNK and NKG2C^+^ and NKG2C^-^ *KLRC1*^-/-^ CAR-iNK cells, to 5 dominant PB-NK cell subsets (**Figure 5D**).^16, 17^ Non-engineered wt iNK cells as well as NKG2A^-^ CAR-iNK cells demonstrated the highest similarity to CD56^bright^ NK cells, supporting the notion that engineering with CAR, CD16 and IL-15/IL-15R drove iNK cell maturation, which is reflected at the phenotypic level by acquisition of NKG2A (**Figure 2A** and **Figure 5D**). NKG2C^+^ and NKG2C^-^ *KLRC1*^-/-^ iNK cells displayed largely overlapping transcriptomes with a slight trend for increased maturation in NKG2C^+^ *KLRC1^-/-^* iNK cells (**Figure 5D**). Indeed, NKG2C^+^ *KLRC1^-/-^* iNK cells expressed higher transcripts of *CD226* (DNAM1), *KLRK1* (NKG2D) and *CD244* (2B4) (**Figure 5E**). This was supported by flow cytometric analysis showing higher levels of cytotoxic mediators (e.g., Granzyme B) and transcription factors (T-bet, Eomes) in NKG2C^+^ *KLRC1^-/-^*iNK cells (**Figure 5F**). Extended CyTOF phenotyping was performed corroborating that NKG2C^+^ *KLRC1^-/-^* iNK cells have slightly higher expression of activating immune receptors, including DNAM-1 NKG2D, 2B4 and NKp30 (**Supplemental Figure 4E**). Together, these findings suggest that *KLRC1* knockout in iNK cells not only triggers compensatory NKG2C surface expression but also enhances their activation and cytotoxic potential through upregulation of key activating receptors and effector molecules.

### Enhanced cytotoxicity of iNK-CAR19 *KLRC1*^-/-^ cells against HLA-E-high targets

Next, we addressed whether the induced surface expression of the NKG2C receptor had any impact on the functional response of CAR19-iNK cells. To this end, we evaluated the CD107a and IFN-γ responses of CAR19-iNK *KLRC1*^-/-^ cells against target cells with varying HLA-E expression levels (**Figure 6A-B**). We found that the natural cytotoxicity response to K562wt was similar for NKG2C^+^ and NKG2C^-^ iNK cells with a slightly better IFN-*γ* production from NKG2C^+^ NK cells (**Figure 6A**). This response was only marginally influenced by the overexpression of HLA-E, suggesting that the main benefit achieved by the *KLRC1* knockout was the reversal of NKG2A-mediated inhibition (**Figure 4C-D**), with a small contribution from spontaneously surfaced NKG2C receptors. Similar results were observed for Nalm-6 lines expressing no, wt, or high levels of HLA-E (**Figure 6B**), although the NKG2C receptor provided a more robust additive signaling to the CAR19 construct against Nalm-6 cells expressing high levels of HLA-E. These results at the subset level were corroborated in assays analyzing the global response of bulk CAR19-iNK *KLRC1*^-/-^ cells against the different targets (**Figure 6C**).

**Figure 6.**
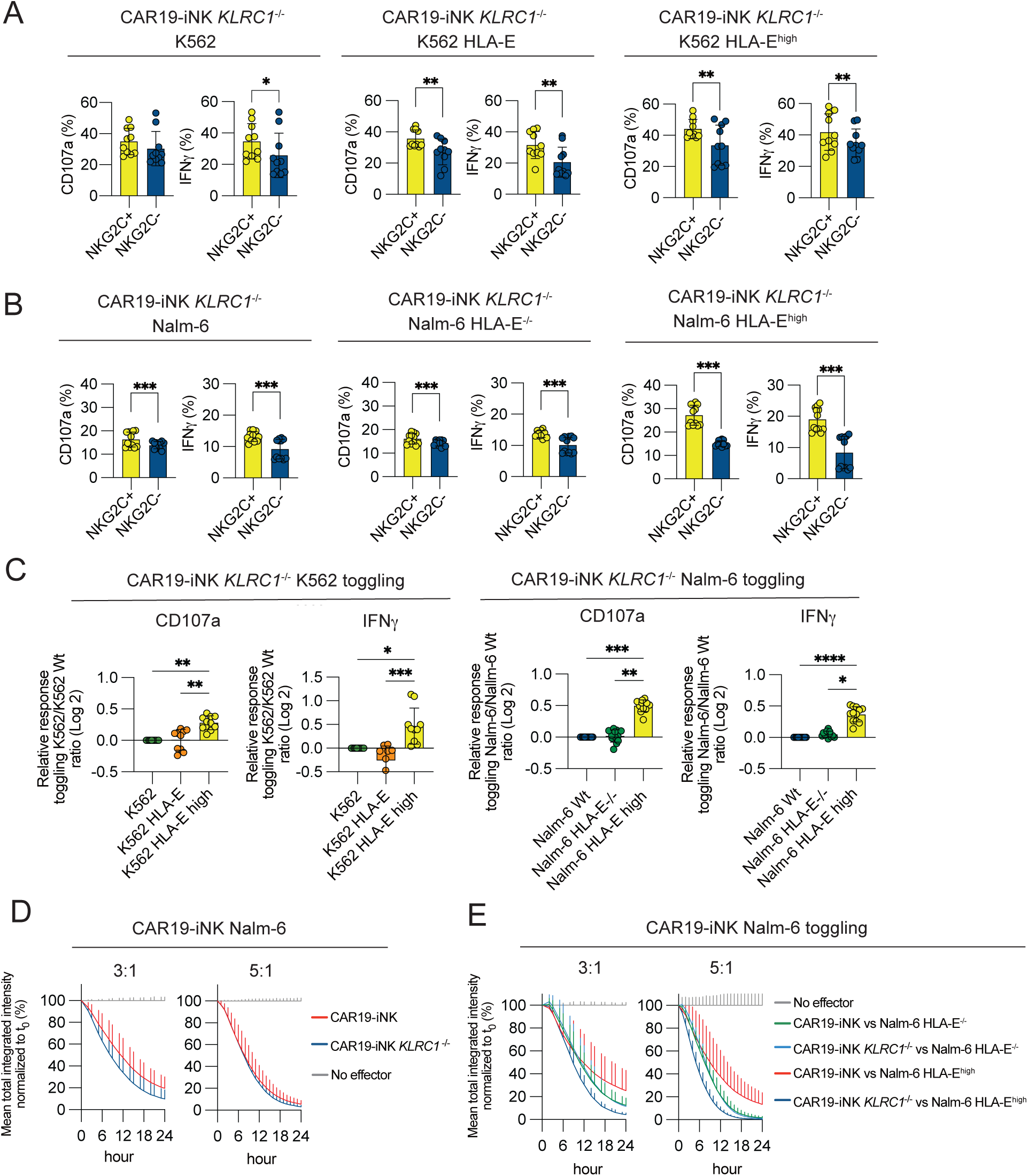
Enhanced cytotoxicity of CAR19-iNK *KLRC1*^-/-^ cells against HLA-E-high targets. (**A-B**) Degranulation (CD107a) and IFNγ production assays comparing CAR19-iNK *KLRC1*^-/-^ cells to wild-type CAR19-iNK cells against K562 (**A**) and Nalm-6 cells (**B**) with varying HLA-E levels. (**C**) Relative response analyses of *KLRC1* knockout cells against target cells with varying HLA-E expression. (**D-E**) Long-term killing assays of CAR19-iNK variants cells against Nalm-6 target cells expressing varying levels of HLA-E. Data are represented as mean (SD). Significance was calculated using a Wilcoxon matched-pairs signed rank test (**A-B**) or a Friedman test followed by Dunn’s multiple comparison test (**C**). p values: * < 0.05, ** < 0.01, *** < 0.001, **** < 0.0001. n = 5-12.

To further evaluate the net effect of the *KLRC1* knockout and the spontaneous surfacing of NKG2C, we performed two series of long-term killing assays. We found that the *KLRC1*^-/-^ iNK cells displayed intact killing responses against Nalm-6wt cells with intermediate levels of HLA-E, corroborating the notion that the NKG2A receptor is not required for the functional education/maturation of iNK cells during their differentiation (**Figure 6D**). Using Nalm-6 variants we observed a titrated response with the lowest killing activity by unedited CAR19-iNK against Nalm-6 HLA-E high cells. We observed intermediate killing activity of unedited CAR19-iNK against Nalm-6 HLA-E knockout cells that was similar to that of *KLRC1*^-/-^ CAR19-iNK against HLA-E knockout cells (**Figure 6E**). In principle, the two latter scenarios reflect the same receptor-ligand relationship, since there is no signal from HLA-E in the Nalm-6 HLA-E knockout target and therefore no beneficial effect of the *KLRC1*knockout. Finally, the strongest killing was observed by *KLRC1*^-/-^ CAR19-iNK against Nalm-6 HLA-E high cells where the added benefit from the surfaced NKG2C receptor comes into play (**Figure 6E**). These findings demonstrate that *KLRC1* knockout phenocopies an adaptive-like state in iNK-CAR19 cells, thereby shifting their functionality toward an HLA-E-responsive profile with enhanced IFN-*γ* production and cytotoxicity. This approach may mitigate the inhibitory effects of NKG2A in immune checkpoint contexts, providing a promising strategy for targeting HLA-E-overexpressing tumors.

## Discussion

The development of off-the-shelf immunotherapies has driven interest in generating NK cells from cord blood stem cells or iPSCs. iPSCs can be derived from accessible sources like skin or peripheral blood and expanded indefinitely *in vitro* without losing pluripotency, ^9^ providing an unlimited supply for cell therapy.^8^ Additionally, iPSCs serve as a versatile platform for multiplexed genetic engineering to enhance specificity, persistence, tumor homing, and functionality of the derived NK cell product.^1, 46^ In this study, we investigated how transcriptional maturation and education through inhibitory receptors and their ligands shape the functional template of iNK cells, a critical foundation for optimizing future engineering strategies.

As a point of departure, we mapped the transcriptomes of a series of iNK lines, either wt (non-engineered) or iNK cells with multiple edits including CAR19, CD16 and tonic cytokine signaling through IL-15/IL-15R complex. We found that iNK cells are classified as bona fide NK cells by CellTypist, a machine-learning-based tool for rapid and automated cell type annotation in single-cell RNA-seq datasets.^34^ Reference mapping placed iNK cells between resting CD56^bright^ and early NKG2A^+^ CD56^dim^ PB NK cells, albeit their signature was by far more similar to IL-15 activated NKG2A^+^ CD56^dim^ NK cells or rapidly cycling KIR^+^ CD56^dim^ NK cells. Thus, iNK cells occupy an intermediate maturation stage with overlapping regulons of CD56^bright^ and CD56^dim^ NK cells, displaying WNT signaling and some shared programs with IL-15-stimulated PB-NK cells (e.g., *MYC*, *LEF*) while also showing unique features like high *EFHD2*, *ALOX5AP*, low *CST7*, *FGFBP2*, *SPON2*, *LYAR*, *SYNE2*, *TIGIT*, and *FCGR3A*, which together suggest they have a relatively “young” transcriptional profile but with a mature cytotoxic machinery.

A hallmark of early NK cell development is the acquisition of inhibitory CD94/NKG2A receptors, which are uniformly expressed on CD56^bright^ NK cells and early CD56^dim^ NK cells.^47, 48^ NKG2A is then gradually lost and its expression inversely correlates with KIRs, which are acquired during late NK cell differentiation, independently of the presence or absence of cognate HLA class I ligands.^17, 49, 50^ In some individuals, infection with CMV drives expansion of adaptive NK cells, a transcriptionally and epigenetically defined terminally differentiated state.^18, 19^ Adaptive NK cells express (oligo)-clonal KIR repertoires and typically, albeit not always, the activating NKG2C receptor.^18, 19, 22^ We found that NKG2A acquisition was more pronounced in iNK lines incorporating multiple gene edits, including tonic cytokine support through IL-15/IL-15R complexes. Although KIR expression was low in all iNK cell lines, it was slightly higher on NKG2A^+^ iNK cells, supporting the notion that NKG2A acquisition defines a population that has reached a more mature stage of differentiation. Indeed, CITE-Seq analysis and reference mapping revealed that NKG2A^+^ iNK cells were most similar to intermediate stages of CD56^dim^ NK cells, while NKG2A^-^ iNK cells matched CD56^bright^ NK cells. We could not identify any iNK cell population that had reached a late or adaptive differentiation stage, characterized by loss of NKG2A and dominance of KIR, and eventually expression of CD57 and NKG2C.

Functional analysis revealed that NKG2A^+^ iNK cells were more responsive to target cell stimulation compared to NKG2A^-^ iNK cells. This suggested a potential role of NKG2A/HLA-E mediated education which tunes the function of PB-NK cells.^24, 26^ To address this possibility, we generated two independent *B2M*^-/-^ iNK lines with no HLA-class I and no HLA-E expression. These lines developed normally and upon acquisition of NKG2A, this subset displayed functional superiority compared to NKG2A^-^ *B2M*^-/-^ iNK cells. These experiments demonstrated that iNK cells develop their functional maturity independently of education. This finding is perhaps not so surprising given that education, or rather, the lack of education, can be overridden by short-term culture in IL-2 or IL-15.^51, 52^ Thus, the priming from long-term culture with feeder cells, genetic edits of IL-15/IL-15R complexes and the addition of exogenous cytokines together drives a functional state that is primarily determined by the stage of differentiation and less by the fine tuning from inhibitory receptors in iNK cell cultures.

HLA-E is emerging as a very prominent checkpoint in immune-oncology.^53, 54, 55, 56, 57^ CRISPR screens in both mouse and humans have identified the Qa-1/NKG2A and HLA-E/NKG2A inhibitory axis, respectively, as a dominant resistant pathway for effective T and NK cell immunotherapy.^58, 59, 60^ The anti-NKG2A mAb Monalizumab is currently in Phase III trials in combination with Durvalumab (anti-PD-L1), a conventional checkpoint molecule, for advanced (Stage III), unresectable NSCLC and in Phase II against non-muscle invasive bladder cancer in patients unresponsive to Bacillus Calmette-Guerin or with exposed cancer in situ. Although HLA-E is ubiquitously expressed by most cells, its expression is induced under inflammatory conditions and has elevated expression in many human tumors.^61, 62, 63, 64, 56, 65, 66, 67, 68^ Moreover, signaling through HLA-E/NKG2A is modulated by the peptide content and is shaped by IFN-*γ*, leading to a more potent inhibitory signal.^69^ Therefore, we next explored to what extent the frequent NKG2A expression on CAR-iNK cells, naturally associated with successful maturation, dampened responses to target cells expressing HLA-E. We found that the inhibitory response to physiological levels of HLA-E was weak and slightly attenuated in iNK cells compared to that of resting PB-NK cells, in line with their primed functional state. However, forced expression of high levels of HLA-E in a single-chain format with β2m and the HLA-G leader peptide VMAPRTLFL resulted in a significant inhibition of degranulation and IFN-*γ* response to natural ligands on K562 targets and CAR19 recognition of Nalm-6 cells. Hence, in situations where there is high expression of HLA-E, possibly induced by IFN-*γ* in the tumor microenvironment (TME),^53^ this interaction may become relevant and constitute an obstacle for successful iNK cell based therapies.

The challenge of high HLA-E expression in the TME has been acknowledged by several research teams and multiple strategies have been employed to overcome this inhibitory checkpoint, including CRISPR editing^70, 71^ and protein expression blockers retaining NKG2A in the endoplasmic reticulum, all of which demonstrated increased targeting of HLA-E high tumors/leukemias by the engineered NK cells.^64^ Here, we silenced NKG2A at the iPSC stage in two independent lines and derived mature iNK cells completely void of NKG2A expression. Our initial worry that these lines would not be functional due to lack of education was partly refuted already by the observation that NKG2A^+^ iNK cells were fully functional in *Β2μ*^-/-^ iNK cells lacking the HLA-E ligand. Indeed, *KLRC1*^-/-^ iNK cells remained responsive to K562wt targets but also against target cells expressing forced high expression of HLA-E. Intriguingly, genetic silencing of *KLRC1* led to spontaneous surface expression of NKG2C. Such compensatory expression has not been previously reported and was not seen in *KLRC1*^-/-^ PB-NK cells. We found that this was not due to aberrant transcription of *KLRC2* located nearby *KLRC1* on chromosome 12 (12p13.2) within the natural killer gene complex (NKC) locus. Instead, all iNK cell lines had abundant *KLRC2* transcripts at baseline. We therefore speculate that NKG2C surfaces as a heterodimer through increased access to free CD94 upon *KLRC1* knockout.

NKG2C is normally expressed at high levels on adaptive NK cell, a clonal state expanding in response to CMV infection or reactivation.^19, 72, 73^ Such adaptive NK cells display a unique epigenetic and transcriptional signature.^16, 18, 74, 75^ We therefore mapped the transcriptome of NKG2C^+^ and NKG2C^-^ iNK cells onto a PB-NK cell reference scoring its similarity with the 5 dominating NK cell subsets, representing key stages of NK cell differentiation.^16, 17^ We found that their signatures were largely overlapping, with a trend for NKG2C^+^ iNK cells to be more similar to intermediate stages of PB-NK cell differentiation expressing slightly higher levels of DNAM-1, NKG2D, KIR and granzyme B but with no evidence of adaptive reprogramming. This supports the notion that the surface expression of NKG2C is not transcriptionally based but rather a consequence of increased access to CD94. Importantly, NKG2C receptors were functional on *KLRC1*^-/-^ iNK cells leading to an increased responsiveness against target cells expressing high levels of HLA-E, surpassing the NKG2C-negative population and acting in cooperation with CAR signaling. Hence, genetic ablation of *KLRC1* in iNK cells may have a double beneficial effect, in not only abolishing potential negative signaling, but also delivering a positive signal.

This work establishes the foundation for engineering iNK cells with distinct receptor profiles to optimize functionality in diverse immunological landscapes. Future investigations should focus on translating these findings into clinical settings, with an emphasis on multiplexed genetic edits to further refine effector function and persistence.

## Methods

### KEY RESOURCE TABLE

Reagents used are listed in the key resource table.

### Derivation of NK cells from human iPSCs

Human iPSCs were differentiated into hematopoietic progenitors and then into CD34+ cells as previously described.^8, 9, 10, 76^ Upon the appearance of CD34+ cells, CD34+ hematopoietic progenitors were transferred into NK cell differentiation B0 medium containing a 2:1 mixture of Dulbecco modified Eagle medium/Ham F12 (Thermo Fisher Scientific, Waltham, MA, 11965092, 11765054), 2mM L-glutamine (Thermo Fisher Scientific, Waltham, MA, 25030081), 1% penicillin/streptomycin (Thermo Fisher Scientific, Waltham, MA, 25030081), 25uM b-mercaptoethanol (Gibco, M3148), 10% heat-inactivated human serum AB (Sigma, H3667-100M), 5ng/ml sodium selenite (Merck Millipore, Burlington, MA, S5261), 50uM ethanolamine (Sigma-Aldrich 02400), 20mg/ml ascorbic acid (MerckMillipore, Burlington, MA, A4544), interleukin-3 (IL-3; R&D Systems Minneapolis, MN, 203-IL); for first week only), stem cell factor (SCF; R&D Systems Minneapolis, MN, 7466-SC), interleukin-15 (IL-15; R&D Systems Minneapolis, MN, 247-ILB), Fml-like tyrosine kinase 3 ligand (FLT3L; R&D Systems Minneapolis, MN, 207-IL). The cells were left in these conditions for 20 days receiving weekly media changes until they had developed into CD45+CD56+CD33-CD3-cells as determined by flow cytometry.

### Cells and cell lines

#### Primary cells

Buffy coats were obtained from anonymous healthy donors from Oslo University Hospital as approved by the regional ethical committee (REK 2018/2485 and 2018/2482). PBMCs were separated by centrifugation using a density gradient medium (Lymphoprep, Serumwerk). Primary NK cells were isolated from PBMCs using the NK cell selection kit (Miltenyi) and according to manufacturer’s protocol. Cells were maintained in RPMI media (Gibco, ThermoFisher) supplemented with 10% FCS (Sigma-Aldrich) and PenStrep (Sigma-Aldrich) at 37°C, 5% CO2.

#### Cell lines

K562 cells, a chronic myeloid leukemia cell line, and Nalm6 cells, a B cell acute lymphoblastic leukemia cell line, were purchased from ATCC. K562 variants and Nalm6 variants were maintained in RPMI media (ThermoFisher) supplemented with 10% FCS (Sigma-Aldrich) and PenStrep (Sigma-Aldrich) at 37°C, 5% CO2. Nalm6^HLA-E-/-^ cells were generated using CRISPR/Cas9 editing. Wild-type K562 as well as Nalm6^HLA-E-/-^ cells were used to generate K562^GspHLA-E^, and Nalm6^GspHLA-E^ by overexpressing HLA-E*01:01 as a single chain construct covalently linked to b2m and to the HLA-G leader sequence peptide VMAPRTLFL (Gsp), as described previously described ^77^.

#### Expansion of PBMC-NK cells and iNK cells

PBMC-NK cells and iNK cells were expanded using the irradiated K562 cells with transduced membrane-bound IL-21 and 4-1BBL in supplemented B0 media for 2 weeks.^78^ K562 cells were propagated in RPMI 1640 media (Sigma) containing 10% fetal bovine serum (FBS; Sigma).

### Flow cytometry assays

Cells were stained following standard flow cytometry procedures. Briefly, cells were collected in a V-bottom 96-well plate (Falcon) and resuspended in 45 μl of FACS buffer (0.5% BSA, 10% FBS in PBS) containing antibodies for extracellular markers. Cells were stained for 30 minutes at room temperature in the dark. For assays that required fixation, cells were fixed using Cytofix/Cytoperm (BD) following manufacturer’s protocol. Full list of antibodies is available in Key Resource Table. To determine HLA-E expression in target cell lines, cells were incubated with anti-HLA-E antibody and a Live/Dead Aqua dye (ThermoFisher) for 30 minutes at 4°C. Samples were then fixed and acquired on flow cytometer.

### Mass cytometry

For viability assessment, cells were stained with Cell-ID Intercalator-103Rh (Fluidigm, San Francisco, CA, 201103B) in complete medium for 20 minutes at 37°C. Maxpar Cell Staining Buffer (Fluidigm, 201068) was used for all antibody staining and subsequent washing. Samples were incubated with Fc receptor binding inhibitor (Thermo Fisher Scientific, 14-9161-73) for 10 minutes at room temperature, before adding surface antibodies and incubating for 30 minutes at 4°C. Subsequently, cells were fixed in Maxpar PBS (Fluidigm, 201058) with 2% paraformaldehyde, transferred to methanol and stored at -20°C. The day after, cells were stained with an intracellular antibody cocktail for 40 minutes at 4°C and labeled with Cell-ID Intercalator-Ir (Fluidigm, 201192B). Samples were supplemented with EQ Four Element Calibration Beads (Fluidigm, 201078) and acquired on a CyTOF 2 (Fluidigm) equipped with a SuperSampler (Victorian Airship, Alamo, CA) at an event rate of <500/sec. Antibodies were either obtained pre-labeled from Fluidigm or conjugated with metal isotopes using Maxpar X8 antibody labeling kits (Fluidigm) (Key Resource Table). FCS files were normalized using Helios software (Fluidigm) and gated on CD45+ CD19- CD14- CD32- CD3- viable single cells using Cytobank (Cytobank Inc., Santa Clara, CA). For subsequent analysis, data was imported into R (R Core Team, 2019) using the *flowCore* package, and transformed using *arcsinh(x/5)*.

### Functional assays

Functional assays were performed at 37°C in RPMI (ThermoFisher) supplemented with 10% FCS (Sigma-Aldrich) and PenStrep (Sigma-Aldrich). Effector cells were incubated with K562 and Nalm6 variants for 4 hours at a ratio of 5:1 with addition of Brefeldin A (GolgiPlug, BD Biosciences). Next, cells were centrifuged at 300 x *g*, surface stained with lineage markers and anti-CD107a antibodies for assessing degranulation. Cells were then fixed with Cytofix/Cytoperm (BD Biosciences) and stained for intracellular cytokines (IFN-*γ*, TNF-α).

### FACS sorting

iNK cells were stained for FACS using the following panel: CD56-ECD, CD45-FITC, NKG2C- PE, NKG2A-APC, CD3-APC/Cy7, CD14-APC/Cy7, CD19-APC/Cy7 and LIVEDEAD Fixable Near-IR Dead Cell Stain Kit, 633 or 635 nm excitation (Invitrogen, L1011922). Cells were sorted using a SONY SH800 Cell Sorter (SONY).

### Incucyte killing assay

Specific tumor lysis was measured in real-time using the Incucyte S3 platform as described previously.^79^ Briefly, target cells stably expressing NucLight Red (Essen Biosciences) were plated and rested overnight and subsequently co-cultured with effector cells at different E:T ratios. Images (3/well) from at least three technical replicates for each condition were acquired every 90 min for 48 hours, using a ×10 objective lens and analyzed by Incucyte v2020A (Essen Biosciences). Graphed readouts represent percentage live target cells (based on NLR expression).

### gRNA, Cas9 protein and RNP formation

CRISPR/Cas9 editing was used to disrupt *HLA-E* in Nalm6 and *KLRC1* in iPSCs. Nalm6 cells were resuspended in SF buffer (Lonza) and mixed with HLA-E RNP or NKG2A RNP in a total volume of 100ul. Following electroporation, Nalm6 were incubated and expanded in their respective medium as described above. HLA-E gRNA target sequence:

5’-ACAACGACGCCGCGAGTCCG

iPSCs: NKG2A gRNAs target sequences:

5’-AAGCTTCTCAGGATTTTCAA, 5’-AGGCAGCAACGAAAACCTAA,

5’-ACTGCAGAGATGGATAACCA

### CITE-seq analysis

Cells were processed following the recommended protocol with the Chromium Single Cell 5’ Library Construction kit (Single Cell 5’ PE Chemistry). The cells were stained using TotalSeq C antibodies, two hashtagging antibodies and 13 surface markers. Both transcriptome FASTQ files and ADT and HTO FASTQ files were processed using the Cell Ranger (v6.1.2) count pipeline. Reads were mapped to the hg38 reference genome. CITE-seq data was analyzed using Scanpy^80^. Quality control was performed by filtering out cells based on number of genes and reads in each cell (minimum 100 genes, maximum 30,000 total counts) and genes were filtered based on number of cells in which the genes were expressed (minimum 5 cells). Cells with more than 10% mitochondrial reads were also filtered out. scVI and scANVI, as implemented in scvi-tools^81^, were used for integration of scRNA-seq data. totalVI^82^ was used for joint modeling of both modalities from the CITE-seq data. Reference mapping was performed using scArches^83^ based on trained models. DEGs were assessed using the differential expression methods built into scVI/scANVI. We performed dimensionality reduction for visualization of the cells by Uniform Manifold Approximation and Projection (UMAP) embeddings based on the embeddings learned from scVI and scANVI. PAGA^35^ was used to quantify the connectivity and to visualize the relationships between different populations of cells. Activity of various gene programs was assessed by AUCell^36^.

### Statistical analysis

Statistical significance was calculated using a Wilcoxon matched-pairs signed rank test, a Friedman test followed by Dunn’s multiple comparison test when comparing more than 2 groups with matched samples, or a Kruskal-Wallis test when comparing more than 2 groups. p values: * < 0.05, ** < 0.01, *** < 0.001, **** < 0.0001. The statistical analysis was performed using GraphPad Prism 10 software.

## Acknowledgements

This project has received funding from the European Union’s Horizon 2020 research and innovation programme under the Marie Sklodowska-Curie grant agreement No. 8382909 (to QH). This work was supported by the Felix Mindus contribution to Leukemia Research (2023-02605) and the Swedish Research Council (2024-02467 to QH). The work was further supported by the Research Council of Norway (275469, 237579, KJM), the Research Council of Norway through its Centres of Excellence scheme, project number 332727 (KJM), the Norwegian Cancer Society (190386, 223310, KJM), The South-Eastern Norway Regional Health Authority (2021-073, 2024- 053, KJM), the Swedish Cancer Society (24 3917 Pj), the Swedish Children’s Cancer Society (PR2024-0148), Knut and Alice Wallenberg Foundation 2018.0106 (KJM), and the US National Cancer Institute (P01 CA111412 (KJM), P009500901 (KJM). We would like to thank the Flow Cytometry Core Facilities at Oslo University Hospital for technical support including CyTOF running and analysis. We would like to thank Genomics Core Facility at Oslo University Hospital for technical support for RNAseq.

## Author contributions

Conceptualization: M.K., J.P.G., B.V. and K.J.M.

Methodology: M.K., C.P., A.C.P. M.L.S., Q.H., M.V., M.T.W., L.T-R., S.G., D.S.K., H.C., B.G., M.D., H.J.H., E.H.A., M.K.K., T.L., H.N., A. P.

Statistics and bioinformatics: M.K., H.N, and A.P.

Formal analysis: M.K., C.P., H.N.; M.L.S., A.P.

Resources: B.G., L.K., A.C.P., S.Z.K.

Writing – original draft: M.K, H.N., A.P., and K.J.M.

Writing – Review & Editing: All authors.

Supervision: J.P.G., J.S.M., J.P.G., B.V. and K.J.M.

Funding acquisition: KJM.

## Data availability

The reference gene expression data used in this paper is available at NCBI GEO with accession number GSE245690 and raw sequencing data is available at EGA with accession number EGAS50000000014. The raw data for the IL-15 stimulated cells is available at EGA with accession number EGAS00001003946. The CITE-seq data from the iNK cell experiments will also be made available at GEO and EGA.

## Code availability

The code for the transcriptional reference map is available on GitHub at http://github.com/hernet/transcriptional-map-nk. The code for the CITE-seq analysis will be made available at https://github.com/hernet/cite_seq_ink.

---

**Supplemental Figure. 1.**
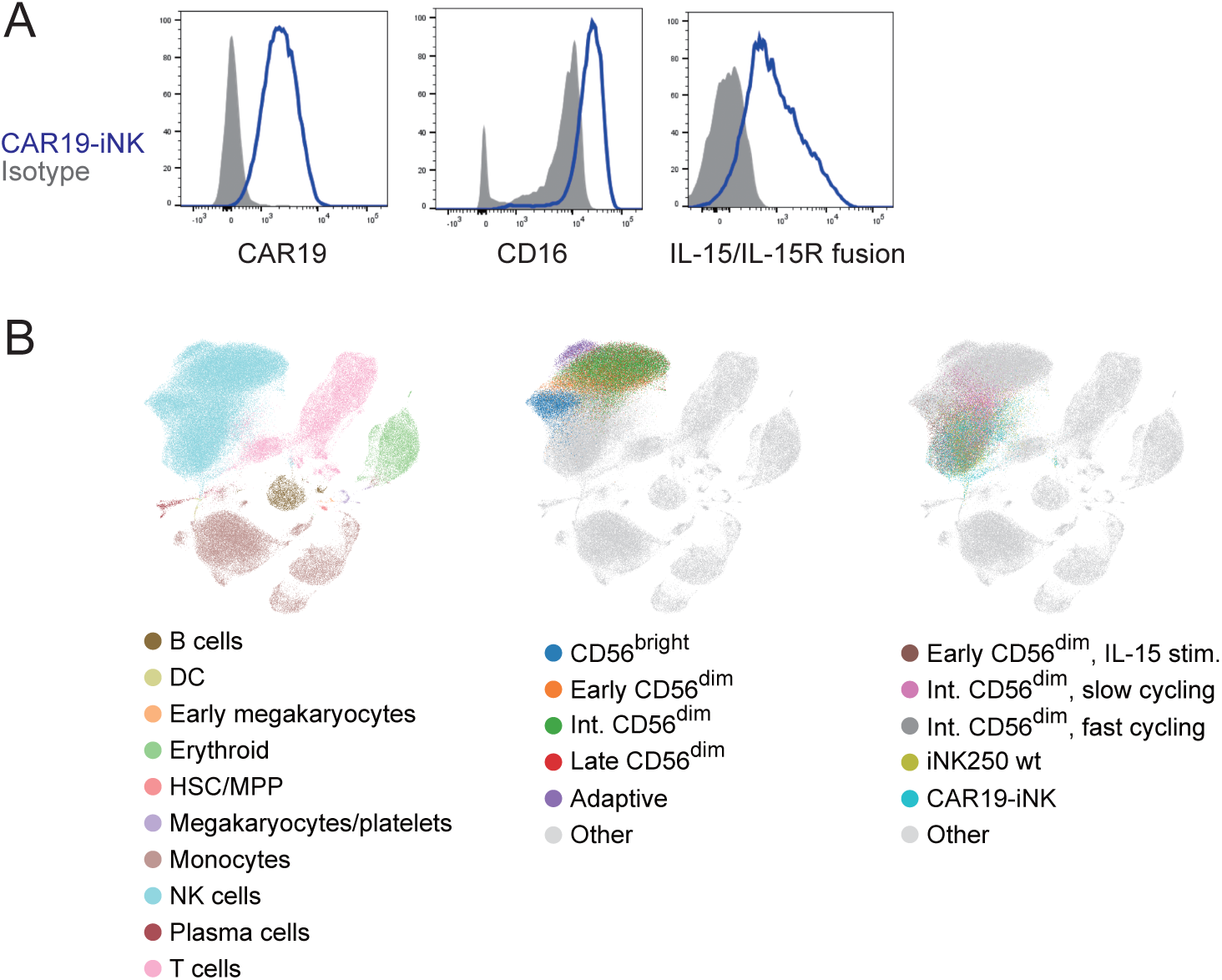
Transcriptional reference mapping of iPSC-derived CAR19-iNK cells. (**A**) Representative FACS plots of CAR19, CD16 and IL-15/IL-15R fusion protein in iNK- CAR19. (**B**) Mapping of iNK cells onto a healthy immune cell reference map. n = 2-12.

**Supplemental Figure. 2.**
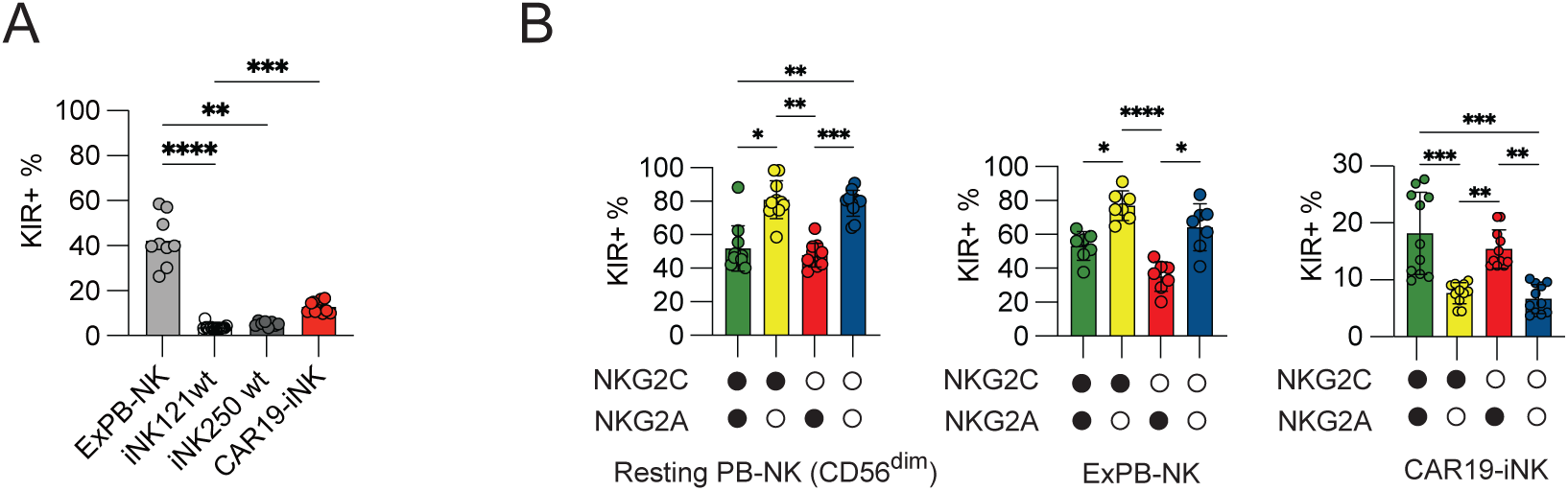
Expression of KIR receptors in subset of iNK cells. (**A**) Percentage of KIR+ subpopulation in each iNK line is shown. KIR^+^ means KIR2DL1 (REA284)^+^ and/or KIR2DL2/3/S2 (GL183)^+^ and/or KIR3DL1 (DX9)^+^. (**B**) Percentage of KIR^+^ population in NKG2C+/- and NKG2A+/- subsets from CD56^dim^ resting PB-NK, expanded PB- NK, and CAR19-iNK. Data are represented as mean (SD). Significance was calculated using a Kruskal-Wallis test (**A**) or a Friedman test followed by Dunn’s multiple comparison test (**B**). p values: * < 0.05, ** < 0.01, *** < 0.001, **** < 0.0001. n = 7-16.

**Supplemental Figure. 3.**
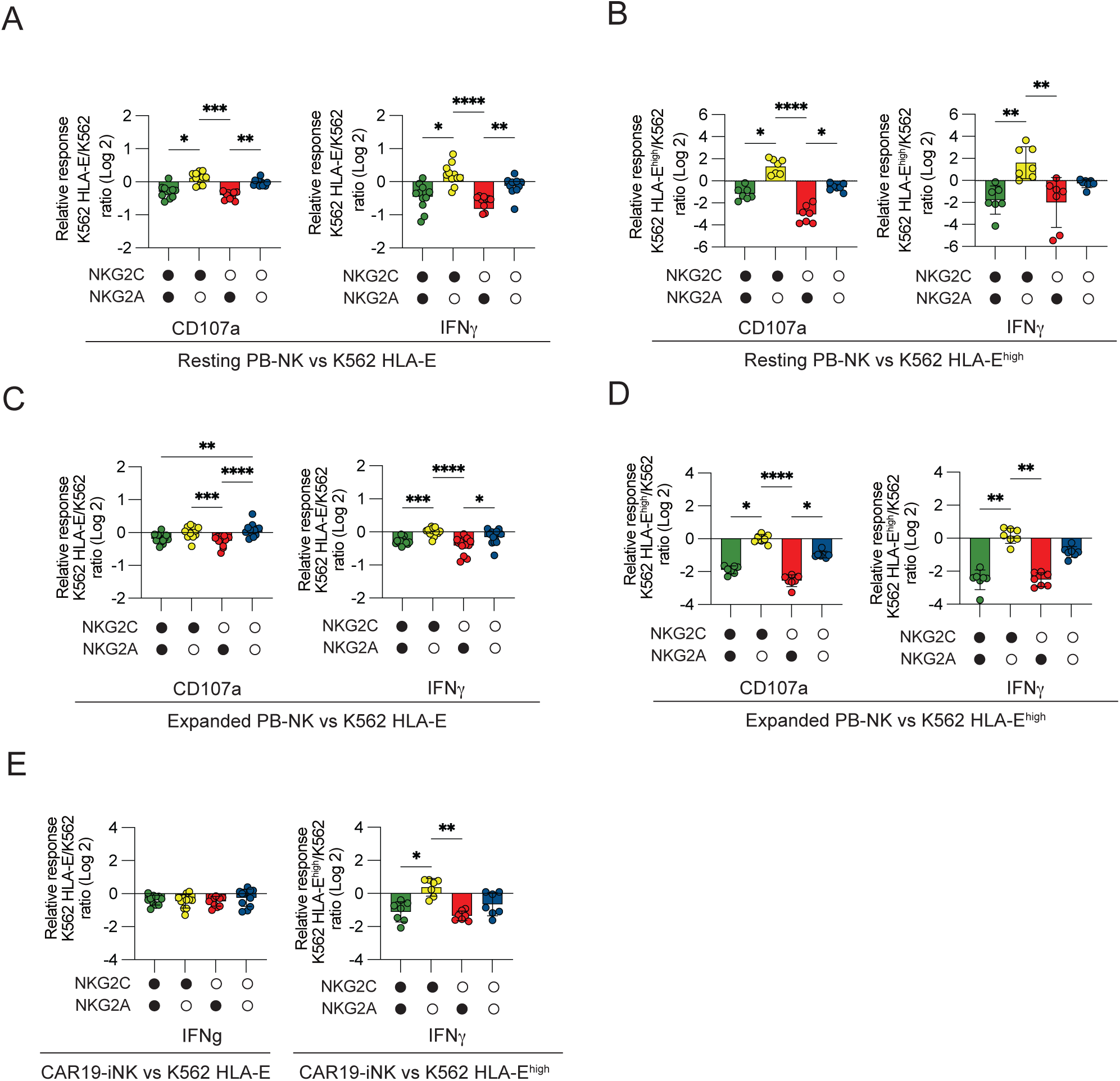
Functionality of iNK cells to HLA-E immune checkpoints on target cells. (**A-E**) Relative response rate of CD107a and IFNγ for K562 HLA-E (**A, C, E**) and K562 HLA-E^high^ (**B, D, E**) in resting PB-NK (**A-B**), expanded PB-NK (**C-D**), and CAR19-iNK cells (**E**) in NKG2C+/- NKG2A+/- subset. Data are represented as mean (SD). Significance was calculated using a Friedman test followed by Dunn’s multiple comparison test. p values: * < 0.05, ** < 0.01, *** < 0.001, **** < 0.0001. n = 7-13.

**Supplemental Figure. 4.**
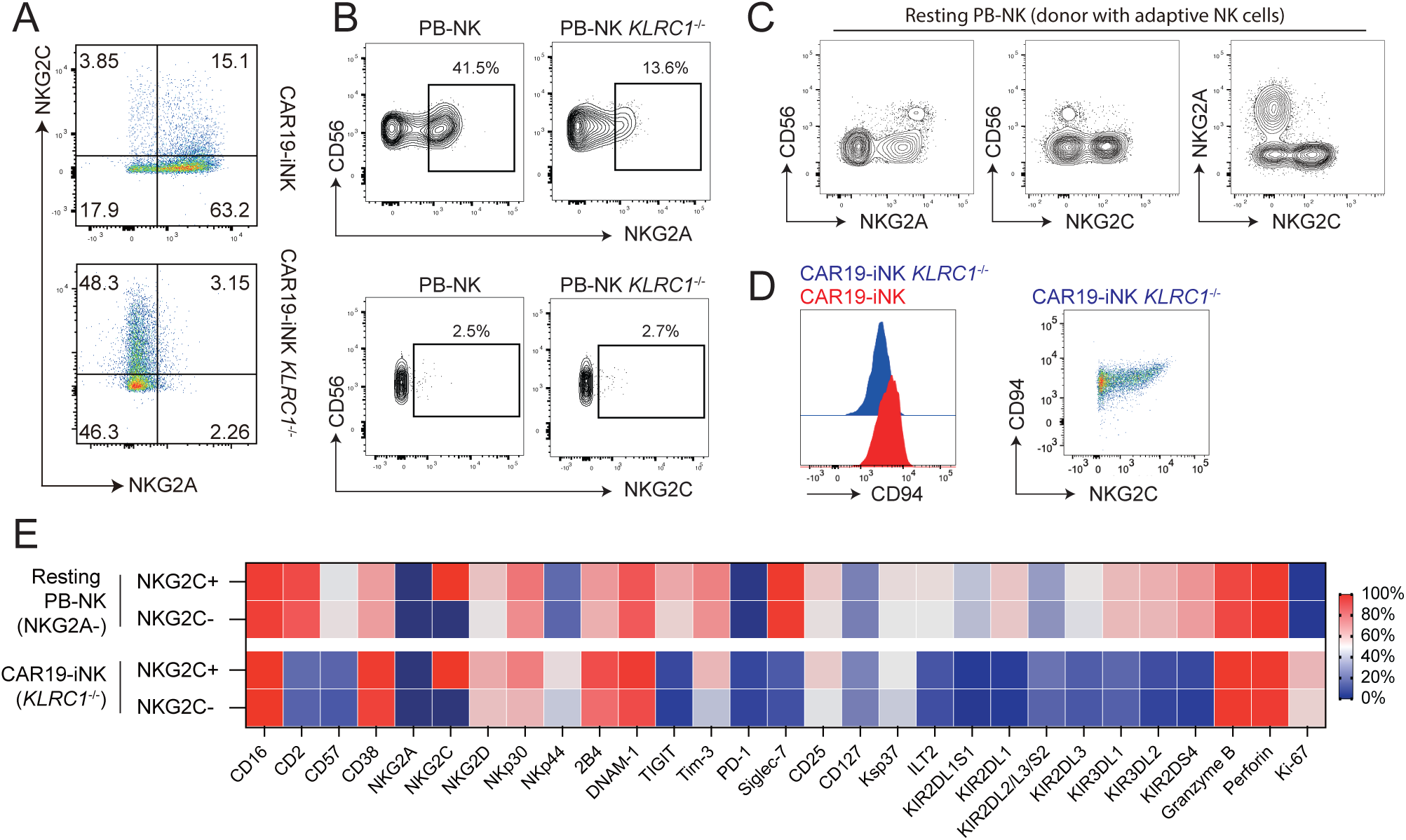
Phenotype of iNK-CAR19 *KLRC1*^-/-^ cells. (**A-C**) Representative FACS histogram of NKG2A and NKG2C expression in CAR19-iNK and CAR19-iNK *KLRC1*^-/-^ cells (**A**), in control and *KLRC1*^-/-^ PB-NK cells (**B**) and in resting PB-NK from a donor with an adaptive NK cell expansion (**C**). (**D**) Representative FACS histogram showing CD94 expression in CAR19-iNK and CAR19-iNK *KLRC1*^-/-^ and co-expression of CD94 and NKG2C in the latter cell line. (**E**) Phenotypic characterization of resting NKG2A- PB-NK and CAR19-iNK *KLRC1*^-/-^ cells, stratified by NKG2C expression. n = 5

